# Space use of invertebrates in terrestrial habitats: phylogenetic, functional and environmental drivers of interspecific variations

**DOI:** 10.1101/2023.07.07.548086

**Authors:** Gwenaëlle Auger, Julien Pottier, Jérôme Mathieu, Franck Jabot

**Affiliations:** UMR 0874 Ecosystème Prairial (UREP), INRAE, Clermont-Ferrand, France; Sorbonne Université, CNRS, IRD, INRAE, Université Paris Est Créteil, Université de Paris Cité, Institute of Ecology and Environmental Sciences of Paris (iEES-Paris), Paris, France

**Keywords:** Space use, allometry, foraging, dispersal, scaling, movement, invertebrates, functional trait

## Abstract

**Aim:** We present the first global database of movement patterns of terrestrial invertebrates, focusing on active dispersal and foraging movements. We depict interspecific variations in movement distances among invertebrates, and assess potential drivers of these variations.

**Location:** Worldwide.

**Methods:** We conducted a meta-analysis using 174 studies from the scientific literature. They provided 401 movement estimates (163 of foraging and 238 of dispersal) from 216 species, 82 families and 22 orders, complemented by the following co-variables: body mass, diet, locomotion mode, tracking method and environmental variables (gross primary productivity and mean temperature of the warmest quarter of the year). We computed allometric relationships between movement distances and body mass both globally and separately for each taxonomic order with sufficient data. We tested the relative influence of the co-variables on movement distances through model selection.

**Results:** We reveal a general positive allometric relationship between movement distance and body mass that holds across most taxonomic orders. We evidence a strong phylogenetic signal in movement distances that translates in variable allometries of movement distances with body mass across taxonomic orders. We further find that interspecific variations of movement distances are primarily driven by functional differences rather than by environmental conditions. Locomotion mode appears to be the most important driver of both dispersal and foraging distances, with larger distances among flying individuals followed by walking and crawling ones for a given body mass. Trophic guild also significantly impacted movement distances with carnivores foraging further than herbivores and decomposers for most body sizes. We finally found little effect of the environmental variables tested.

**Main conclusions:** Our study provides general allometric equations for terrestrial movement distances of invertebrates. It further reveals important functional drivers of their interspecific variation in space use with a dominant role of their evolutionary history.

## INTRODUCTION

Animal movements have widespread consequences at population, community, ecosystem and evolutionary levels. They alter local population density and growth rate through emigration and immigration (Law et al. 2003) as well as metapopulation dynamics and evolution (Hanski and Gaggiotti 2004). At the community level, previous research has mainly focused on the role of dispersal limitation in metacommunity dynamics (Holyoak et al. 2005) and the associated response of biodiversity to climate changes (Lenoir et al. 2020). The key role of other types of movement for community dynamics has also been stressed, in particular the role of foraging movements in the spatial dynamics of foodwebs (Amarasekare 2008) and more generally community assembly (Schlägel et al. 2020). Animal movements also couple the dynamics of distinct habitats by their associated transfer of matter and energy. A substantial body of theory on such meta-ecosystem dynamics has been developed (Gounand et al. 2018; Guichard et Marleau 2021), and available data demonstrate the significance and breadth of such transfers (Gounand et al. 2018; McInturf et al. 2019).

Animals perform different types of movements at various spatial and temporal scales and for a variety of reasons. Four basic movement types are generally considered (Barton et al. 2015). First, dispersal is generally defined as a unidirectional movement leading to gene flow between distinct populations. This process can either be active, like the mechanical flight of most beetles, or passive, like the ballooning of some spiders with wind currents (Bishop 1990) or the phoresy of tiny organisms on larger ones (Bartlow and Agosta 2021). Second, foraging movements are the way animals daily explore their environment for food resources. They are restricted for many animals to a compact area called home range (Burt 1943). Third, nomadism refers to the movement pattern of an animal that irregularly shifts its home range core location to exploit spatially and temporally fluctuating resources. This type of movement is best described in birds and mammals, but also occurs in a range of diverse taxonomic groups including gastropods (Posso et al. 2012) or even social insects that can occasionally relocate their nests (McGlynn 2012). Fourth, migration occurs when an animal seasonally undertakes a bi-directional movement. This type of movement connects separated breeding and non-breeding habitats of migratory vertebrates like amphibians and some large mammals, birds or fish. Migratory insects, like some fly, butterfly or moth species (Hawkes et al. 2022, Chowdhury et al. 2021) also migrate over long distances to deal with seasonal variations of resource availability (Dingle 2014).

Movement ecology has developed rapidly over the last decade with the development of a unified paradigm (Nathan et al. 2008). A general understanding of the drivers and spatial extents of animal movements is indeed of particular relevance for diverse research topics, including ecosystem modeling (Earl and Zollner 2017), biological control (McEvoy 2018) or niche tracking in a context of global changes (González-Varo et al. 2017). To understand the commonalities in movement patterns across the animal kingdom, data syntheses are needed to document the magnitude and variability of movement rates among and within species, but also to understand their drivers. Several syntheses revealed that interspecific variability in movement distances is mostly driven by functional and life-history traits in vertebrates. Among these traits, body size has been found to be the main factor that correlates positively with migration (Hein et al. 2012), foraging (Tamburello et al. 2015) and dispersal distances (Santini et al. 2013). Locomotion mode (flying, walking or swimming) alters the intercept and slope of these allometries between movement and body size because of the varying penetrability of the associated medium (air, land or water). Larger movement distances are observed in more penetrable media (Santini et al. 2013, Tamburello et al. 2015, Straus et al. 2022). These studies also evidenced that diet is a significant determinant of space use with carnivore foraging and dispersing over larger distances than herbivores to compensate for lower resource densities. They have also demonstrated a significant phylogenetic inertia of movement distances with taxon-specific allometric relationships with body size.

Similar syntheses on invertebrate taxa are currently lacking. Although invertebrates represent 75% of all described species on Earth and almost 95% of all animal species (Eisenhauer et Hines 2021), a general picture of the variability of movement distances and their drivers is still lacking for this large group of animals. Hirt et al. (2017) revealed a positive scaling of exploratory speed with body mass across six classes of invertebrates, but they did not study the movement distances of these taxa. We therefore aim at filling this gap and at assessing whether invertebrate movement distances are influenced by the same set of functional traits as vertebrate taxa **(Figure 1a-d)**. We focus on active dispersal and foraging distances, since data about invertebrate nomadism and migration are scarce (Hein et al. 2012). We also contend that animal movements depend on the abiotic environmental context and we aim at assessing such abiotic drivers (McManus 1988). We specifically study the effect of i) temperature because of the ectothermic status of invertebrates (Gibert et al. 2016), and ii) gross primary productivity because it may constrain food resource availability and thus correlate negatively with foraging and dispersal distances (Bastille-Rousseau et al. 2017; Abrahms et al. 2021) **(Figure 1e)**.

**Figure 1.**
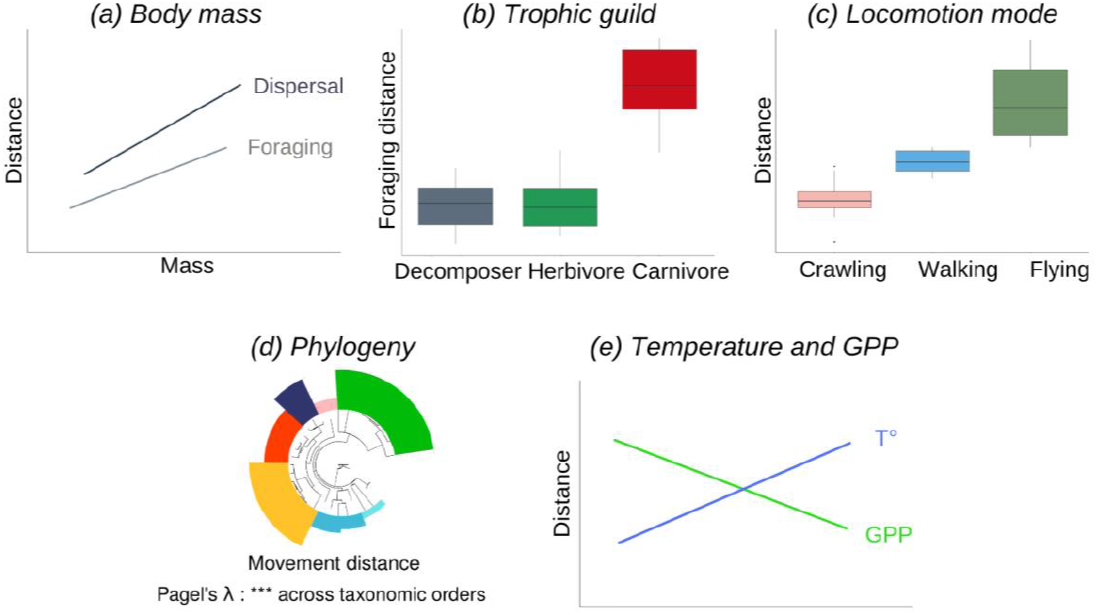
Tested predictions: *(a)* Positive scaling of dispersal and foraging distances with body mass ; *(b)* Carnivores are expected to forage further than herbivores and decomposers ; *(c)* Flying individuals move further than walkers, both moving further than crawlers ; *(d)* Phylogenetic signal in dispersal and foraging movement distances ; *(e)* Higher local temperature leads to larger movement distances. Larger gross primary productivity *(GPP)* leads to lower movement distances.

## METHODS

### Literature search and data selection

We conducted a literature search on the Web of Science and Google Scholar with the title request ((“invertebrate” OR [any known order of terrestrial invertebrate]) AND (“space use” OR “home range” OR “foraging” OR “dispersal” OR “movement pattern” OR MRR OR CMR OR telemetry OR “harmonic radar”)). Specific words relative to aquatic habitats were specified as unwanted, as well as journal categories like “Toxicology” or “Neurosciences” (see full research strategy in Supporting information). We looked for cited references to similar works within each selected paper and added them to our database when relevant. We found three reviews about movement of specific taxonomic groups, and the original papers were double-checked in order to ensure that the concepts of foraging distance, home range and dispersal were used according to consistent definitions. For instance, Mazzi et Dorn (2012) conducted their review by considering the term ‘dispersal’ in a very broad sense, including short distance foraging flights which we incorporated in our database as home range measures (Schlägel et al. 2020).

Studies were retained if they provided quantitative information about mean active dispersal distance (m) and/or mean foraging distance (m) or home-range size (m^2^) for terrestrial invertebrate individuals. We did not compile data of home range size of social insects’ colonies (ants or termites colonies for instance) because they do not correspond to individual traits but to emergent collective patterns. We also excluded genetic and simulation studies that only provided indirect measures of movement distances. Translocation experiments were also excluded, because the ability of an individual to return to its initial territory beyond a certain distance does not only depend on its motion capacities, but also on its memory of visual landmarks and other sorts of cues (Able 1980). The duration of the studies that we retained varied from a single to a few days or even weeks, because all species do not need the same amount of time to complete a dispersal event. We assumed that the retained dispersal studies tracked individuals’ movements during the whole duration of their dispersal event.

### Foraging and dispersal distances

We distinguished dispersal from foraging movement depending on the life stage of the tracked individuals, the duration of the study and the period of the year in which the study took place, which helped us distinguish between movements of natal, post-emergence or breeding dispersal and daily routine movements. We defined dispersal distance as the longest straight-line displacement of an individual between the first and last time it was observed. When necessary, we performed complementary searches to ensure that the studied species was not migratory or not observed during a seasonal migration period. We defined foraging distance as the average maximal distance from the point of release in tracking studies, or from the nest in which individuals were daily observed. When studies provided only a home range size in square meters, we used the mean radius of this area as a proxy for foraging distance.

### Body mass

We reported the mean dry mass of the group of individuals whose movements were studied when available within the publication. Otherwise, we performed a complementary literature search of body mass of the studied taxon. When only fresh mass or body length values could be found, dry mass was calculated thanks to regression coefficients from the literature for distinct taxonomic groups (Sabo et al. 2002; James et al. 2012; Sage 1982; Newton et Proctor 2013; Petersen 1975). When only a mean body length or fresh mass value was available, and when no allometric equation was applicable, we estimated a species’ dry mass with an allometric equation of a phylogenetically close species. Calculation details for the dry mass of each species are available in the **Appendix 1**.

### Locomotion mode

In terrestrial environments, invertebrates actively move in three different ways: they either fly, walk or run (non-alate OR alate species moving on the ground with articulated legs) or crawl (above-ground or below-ground limbless species or larvae). Many invertebrate species undergo a shift in their locomotion mode during their lifetime (e.g. Crawling lepidoptera caterpillars become flying imagos). We therefore associated to each observation in our database the locomotion mode corresponding to the exact life stage of the individuals at the time of the study, and used the corresponding body mass and trophic guild of this life stage.

### Trophic guild

We classified species as either carnivores, herbivores or decomposers. We performed this classification in broad trophic groups to avoid a multiplication of specific trophic habits in our database that would have small sample sizes and for which we would not have clear *a priori* predictions. Hence, we classified omnivorous species feeding on both plants and other invertebrates as carnivores. We grouped hematophagous species like ticks or mosquitoes with carnivores. We pooled granivore and palynivore species with herbivores, and xylophagous, saproxylic, fungivore, detritivore and coprophagous animals as decomposer species.

### Habitat

We distinguished *in situ* studies from *ex situ*, laboratory studies. We extracted the location of the study, or used approximate geographic coordinates based on the description of the area, or the country or state’s centroid for the few studies that did not provide a precise location of their experiment. Two environmental variables were extracted from the location of the studies: the mean temperature of the warmest quarter of the year (*Bio10*, resolution 1×1 km, Fick and Hijmans 2017), related to the species’ ability to get active; and the gross primary productivity (global *GPP* data for year 2016, 5.5 km grid, Zhang et al. 2017), related to the resource abundance of herbivore species under the assumption that resource quality for herbivores is positively correlated with its quantity (Loranger et al. 2014).

### Methodology

Finally, we also reported the methods used to track individuals (capture-mark-recapture, telemetry, harmonic radar, flight mill, visual monitoring or video tracking). The combination between the study condition (*in situ* or *ex situ*) and the tracking method created a new “Method” variable that we used as a random effect in our statistical models (e.g. “*In situ* : CMR” or “*Ex situ* : Flight mill”).

### Statistical analysis

#### Taxon-specific allometry of space use

We first performed ordinary least square (OLS) regressions to assess the allometry of space use for each taxonomic order that presented more than seven observations in our database. Since we found that movement distances estimated by *ex situ* studies are significantly larger than those estimated *in situ* (dispersal : *ANCOVA* : F = 78.161, p = 2e-16; foraging : *ANCOVA* : F = 36.945, p = 8.58e-09), we performed these analyses on ‘*in situ*’ studies only, except for the *Haplotaxida* order for which we only found foraging data from laboratory studies. We log-transformed (Log_10_) body mass and movement distances for these analyses. For each taxonomic order, we performed either mixed models with tracking method incorporated as a random effect on the intercept value, when several types of tracking methods were used, or simple linear models otherwise. We made this choice of mixing linear and mixed models to follow the same methodology as the one of Tamburello et al. (2015) in their analyses of movement of vertebrate data.

#### Functional and environmental drivers of interspecific variations in space use

We first tested the influence of functional drivers on movement distances, using the full dataset of *in situ* and *ex situ* studies. We assessed whether the locomotion mode and trophic guild influenced the coefficients of our initial allometric models “(Dispersal or foraging) Distance ^ Dry Mass”. We built mixed models to test each predictor separately for both dispersal and foraging data sets, with tracking method and taxonomic order incorporated as random effects on the intercept value. These random effects allowed us to deal with multiple tracking methods while accounting for phylogenetic interdependence. We tested interaction effects between body mass and locomotion mode, and between body mass and trophic guild. We finally combined these two hypothesized functional drivers into a “full” model. The relative support of each model was assessed with the Akaike Information Criterion corrected for small samples (AICc). The marginal and conditional R^2^ of the models were calculated to assess the proportion of variance explained respectively by the fixed effects alone, and by both fixed and random effects, with the “r.squaredGLMM” function from MuMIn (Barton 2022).

A second analysis aimed at assessing the relative influences of functional and environmental drivers using the dataset of *in situ* studies only. We built a mixed model with the following potential drivers of space use: body mass, trophic guild, locomotion mode, temperature and GPP. We computed the partial eta-squared of these different predictors to assess their relative influence on movement distances in our dataset, with taxonomic order and tracking method again included in the models as random effects on the intercept value. All analyses were performed on R 4.2.1 (2022.06.23).

### Phylogenetic signal

We also performed a phylogenetic analysis of the interspecific variation in movement distances. We constructed phylogenetic trees with dated branches thanks to the OneTwoTree pipeline (Drori et al. 2018) for a subset of the species in our database (174 out of 216). We calculated the phylogenetic signal in movement distance using Pagel’s lambda metric that is adapted to continuous response variables. Pagel’s λ ranges from 0 if there is no link between the response variable and phylogeny, to 1 if closely related species respond exactly the same way to the predictor variables (Pagel 1997, Pagel 1999). Because phylogenetic relatedness between species is likely to cause an inter-dependency of the observations of movement distances, we computed phylogenetic generalized least squares regressions (PGLS) using the ‘nlme’ package in R (Pinheiro and Bates 2023) to account for interspecific autocorrelation and we compared the PGLS regression results with the fit of non-phylogenetic, ordinary least square regressions (OLS).

## RESULTS

### Datasets

We assembled a database of 174 scientific articles that met our selection criteria. They provided 401 movement observations of individuals from 216 species, 82 families and 22 orders. These data correspond to observations of dispersal (238, 59%) and foraging (163, 41%) distances, with a heterogeneous distribution between trophic and locomotion groups **(see Appendix 3, Table 1 in Supporting information)**. Biogeographical realms are unequally represented, with 80% of the total observations coming from the Nearctic and Palearctic world regions, while only 14% is from the Southern hemisphere of the globe **(Figure 2)**. However, the two environmental variables in our database (temperature and gross primary productivity) still cover a large and continuous spectrum of values **(see Appendix 3, Figure 1 in Supporting information)**.

**Table 1.**
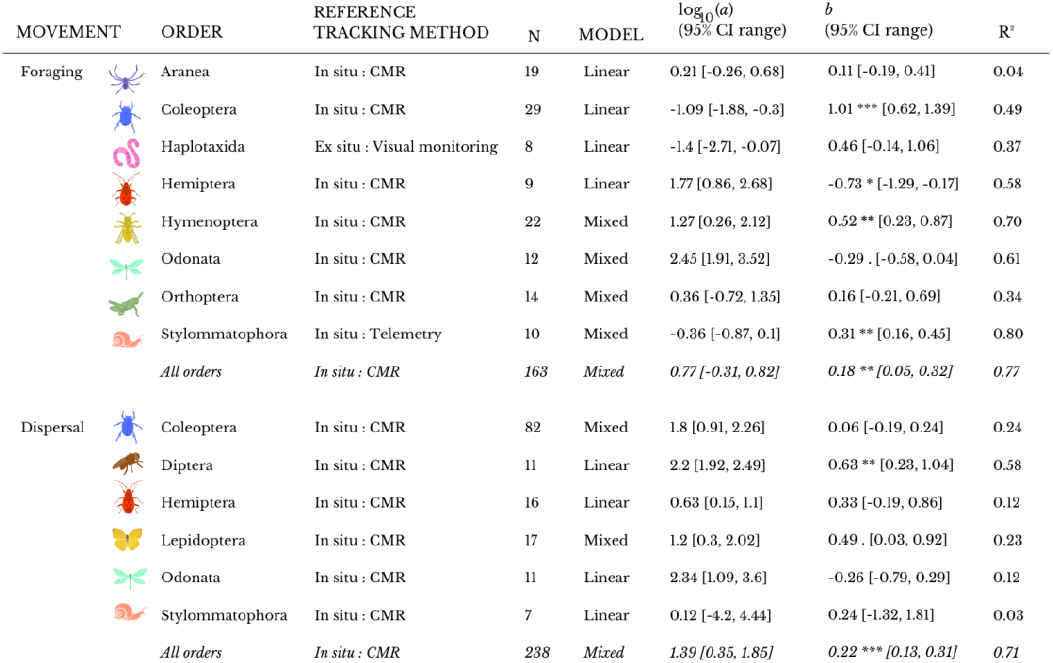
Regression parameter estimates for body mass across separate taxonomic orders with n ≥ 7 observations. Global regressions (*‘All orders’*) include all taxonomic orders in our databases without constraint on minimal number of data. Simple linear models are of the form log_10_(D) ^ log_10_(a) + b x log_10_(DM) where D : movement distance and DM : dry mass. Mixed linear models include the tracking method as a random effect. We used the most frequently used tracking method as the reference for each taxonomic order. log_10_(a): intercept; b: slope; 95% CI: 95% confidence interval. R^2^ corresponds to the conditional R^2^ for mixed models. Significance (p-values) codes : 0 ‘***’ 0.001 ‘**’ 0.01 ‘*’ 0.05 ‘.’ 0.1 ‘ ‘.

| MOVEMENT | ORDER | REFERENCE TRACKING METHOD | N | MODEL | $\log_{10}(a)$<br>(95% CI range) | $b$<br>(95% CI range) | R<sup>2</sup> |
| --- | --- | --- | --- | --- | --- | --- | --- |
| Foraging | Aranea | In situ : CMR | 19 | Linear | 0.21 [-0.26, 0.68] | 0.11 [-0.19, 0.41] | 0.04 |
|  | Coleoptera | In situ : CMR | 29 | Linear | -1.09 [-1.88, -0.3] | 1.01 *** [0.62, 1.39] | 0.49 |
|  | Haplotaxida | Ex situ : Visual monitoring | 8 | Linear | -1.4 [-2.71, -0.07] | 0.46 [-0.14, 1.06] | 0.37 |
|  | Hemiptera | In situ : CMR | 9 | Linear | 1.77 [0.86, 2.68] | -0.73 * [-1.29, -0.17] | 0.58 |
|  | Hymenoptera | In situ : CMR | 22 | Mixed | 1.27 [0.26, 2.12] | 0.52 ** [0.23, 0.87] | 0.70 |
|  | Odonata | In situ : CMR | 12 | Mixed | 2.45 [1.91, 3.52] | -0.29 . [-0.58, 0.04] | 0.61 |
|  | Orthoptera | In situ : CMR | 14 | Mixed | 0.36 [-0.72, 1.35] | 0.16 [-0.21, 0.69] | 0.34 |
|  | Stylommatophora | In situ : Telemetry | 10 | Mixed | -0.36 [-0.87, 0.1] | 0.31 ** [0.16, 0.45] | 0.80 |
|  | *All orders* | *In situ : CMR* | 163 | *Mixed* | *0.77 [-0.31, 0.82]* | *0.18 \*\* [0.05, 0.32]* | *0.77* |
| Dispersal | Coleoptera | In situ : CMR | 82 | Mixed | 1.8 [0.91, 2.26] | 0.06 [-0.19, 0.24] | 0.24 |
|  | Diptera | In situ : CMR | 11 | Linear | 2.2 [1.92, 2.49] | 0.63 ** [0.23, 1.04] | 0.58 |
|  | Hemiptera | In situ : CMR | 16 | Linear | 0.63 [0.15, 1.1] | 0.33 [-0.19, 0.86] | 0.12 |
|  | Lepidoptera | In situ : CMR | 17 | Mixed | 1.2 [0.3, 2.02] | 0.49 . [0.03, 0.92] | 0.23 |
|  | Odonata | In situ : CMR | 11 | Linear | 2.34 [1.09, 3.6] | -0.26 [-0.79, 0.29] | 0.12 |
|  | Stylommatophora | In situ : CMR | 7 | Linear | 0.12 [-4.2, 4.44] | 0.24 [-1.32, 1.81] | 0.03 |
|  | *All orders* | *In situ : CMR* | 238 | *Mixed* | *1.39 [0.35, 1.85]* | *0.22 \*\*\* [0.13, 0.31]* | *0.71* |

**Figure 2.**
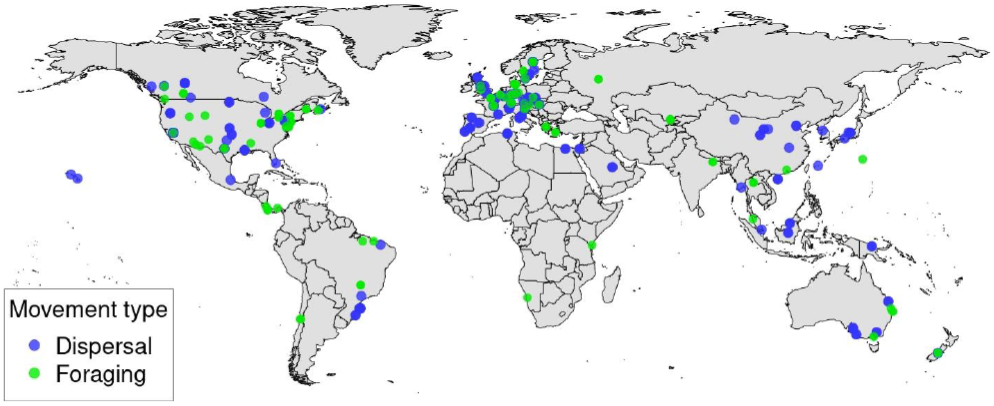
Location map of the data included in the meta-analysis.

Our data compilation encompasses wide ranges of movement distances (from 10^−2^ to 105 m) and of body masses (from 10^−4^ to 104 mg) **(see Appendix 3, Figure 2 in Supporting information)**. Data sources are provided in **Appendix 2**.

### Allometry of space use

Body mass significantly explains both dispersal and foraging distances (OLS regressions: R^2^ = 0.24 and 0.20, respectively), but with a far lower predictive power than previous synthesis studies conducted on vertebrates. The Pagel’s λ-statistic shows strong and highly significant phylogenetic signals for both dispersal and foraging movement distances (Dispersal : λ = 0.99, p = 1e-22, n = 108 ; foraging : λ = 0.89, p = 5e-12, n = 76). As a result, phylogeny slightly improves the fit of the allometric regressions (PGLS : dispersal : R^2^ = 0.26; foraging : R^2^ = 0.21). Also, for a given body mass, dispersal distance is significantly larger than foraging distance (*ANCOVA* : F = 136.74, p < 2.2e-16), which is consistent with the definitions we used for these two movement types.

When looking at the different taxonomic orders separately, we confirm a strong phylogenetic signal in movement distances with a wide variation of regression estimates among taxonomic orders for both foraging **(Figure 3)** and dispersal **(Figure 4)** movements. We recover a positive relationship between body mass and movement distances for most taxonomic orders, but with variable slopes **(Table 1)**. Only *Odonata* individuals have a non-significant negative correlation between body mass and both dispersal and foraging distances and *Hemiptera* individuals have a significant negative correlation between body mass and their foraging distances **(Table 1)**. We also note a strong dispersion of data points of Coleoptera dispersal distances around the regression line, whose slope coefficient is almost null (b = 0.06, **Table 1**).

**Figure 3.**
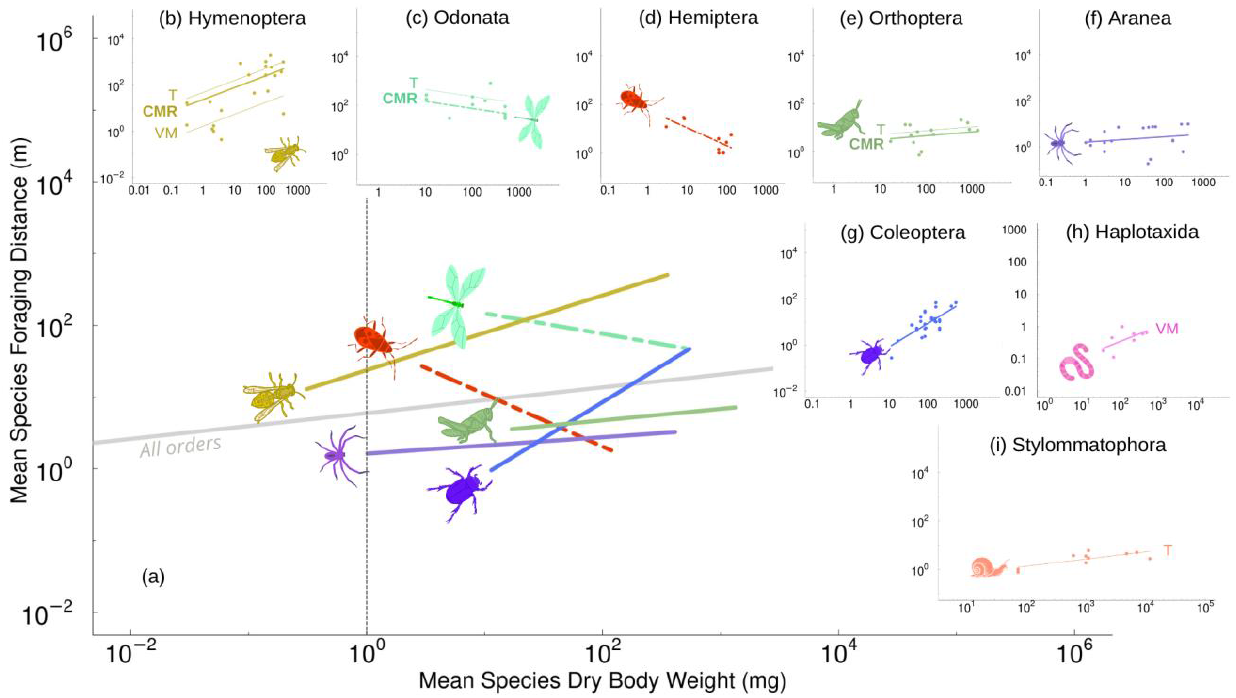
Allometry of foraging distances in terrestrial invertebrates across taxonomic orders (n ≥ 7 obs.). (a) Summary of taxonomic order-specific regression lines of capture-mark-recapture (CMR) data; (b) flying (wasps and bees) and walking (ants) Hymenoptera ; (c) dragonflies and damselflies (Odonata) ; (d) true bugs (Hemiptera) ; (e) grasshoppers, locusts and crickets (Orthoptera) ; (f) spiders (Aranea) ; (g) beetles (Coleoptera) ; (h) annelid worms (Haplotaxida) ; (i) snails and slugs (Stylommatophora). In panels b, c, e and h, regression lines of alternative tracking methods are also reported: T: telemetry ; VM: visual monitoring.

**Figure 4.**
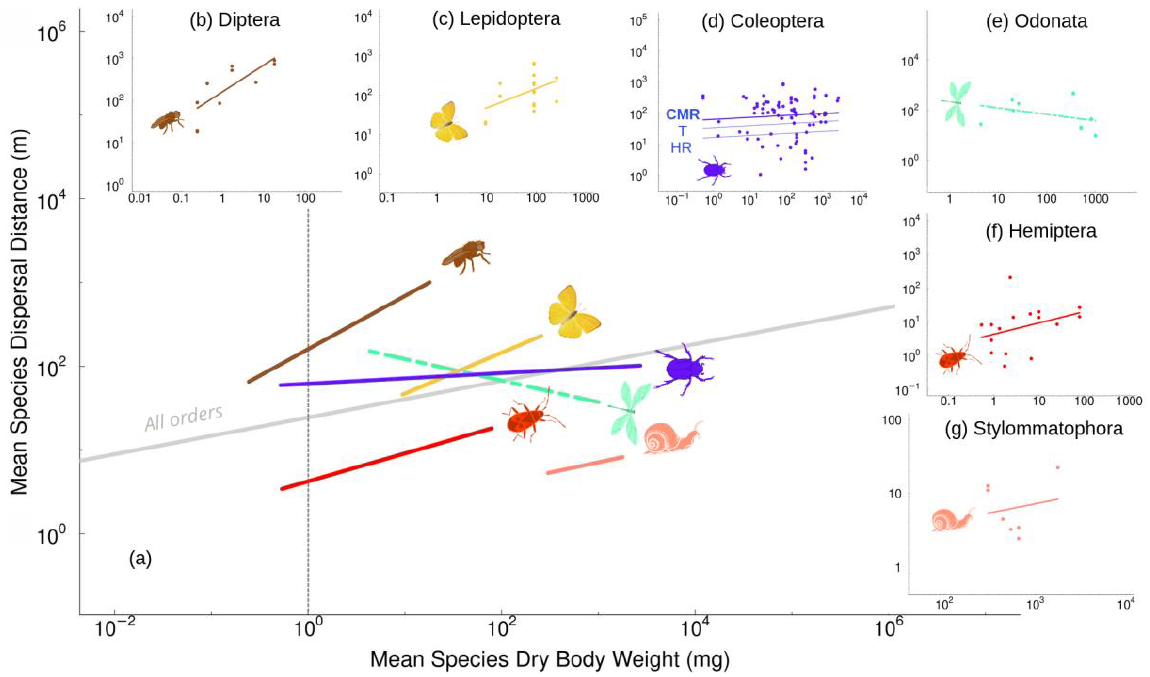
Allometry of dispersal distances in terrestrial invertebrates across taxonomic orders (n ≥ 7 obs.). (a) Summary of taxonomic order-specific regression lines of capture-mark-recapture (CMR) data ; (b) Diptera ; (c) butterflies (Lepidoptera) ; (d) beetles (Coleoptera) ; (e) dragonflies and damselflies (Odonata) ; (f) true bugs (Hemiptera) ; (g) snails and slugs (Stylommatophora). In panel d, regression lines of alternative tracking methods are also reported: T : telemetry ; HR : harmonic radar.

### Effects of intrinsic and environmental factors on movement distances

We found that locomotion mode is the most influential driver of interspecific variations in foraging **(Figure 5a)** and dispersal **(Figure 5b)** distances. The other drivers have comparatively low effects with partial η^2^ values below 0.1. Trophic guild is the second most influential driver, followed by body mass and the two environmental drivers tested (temperature and gross primary productivity, **Figure 5**). For consistency with the other analyses, **Figure 5** includes taxonomic order and tracking method as random effects. However, phylogeny predominates all other factors when tested as a fixed effect **(see Figure 2 in Supporting information)**.

**Figure 5.**
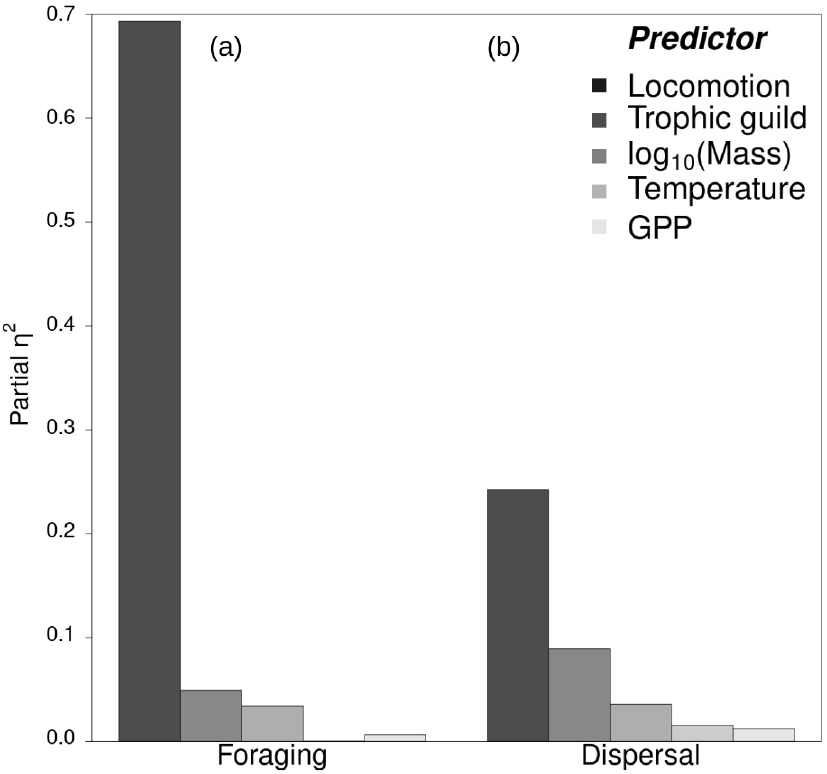
Effect sizes of locomotion mode, body dry mass, diet, temperature and GPP as drivers of invertebrate foraging (a) and dispersal (b) movements. Taxonomic order and tracking method are included in the models as random effects. Note that *in situ* studies only have been considered for this figure to allow the inclusion of the two environmental variables, temperature and gross primary productivity.

We found a significant interaction effect between trophic guild and body mass for foraging (*ANOVA* : F = 7.857, p = 0.0005) and dispersal movements (*ANOVA* : F = 3.037, p = 0.049) with a decrease in AICc in models with interaction of 3.9 and 10.7 respectively **(Table 2)**. Our a priori prediction that carnivores should forage further than the two other trophic guilds was only partially supported concerning foraging movements, except for the largest organisms **(Figure 6a)**. This pattern did not however extend to dispersal movements. We found decomposers to have the highest intercept value followed by carnivores, both groups showing similar allometric regression slopes that are crossed by that of herbivores **(Figure 6b)**.

**Table 2.**
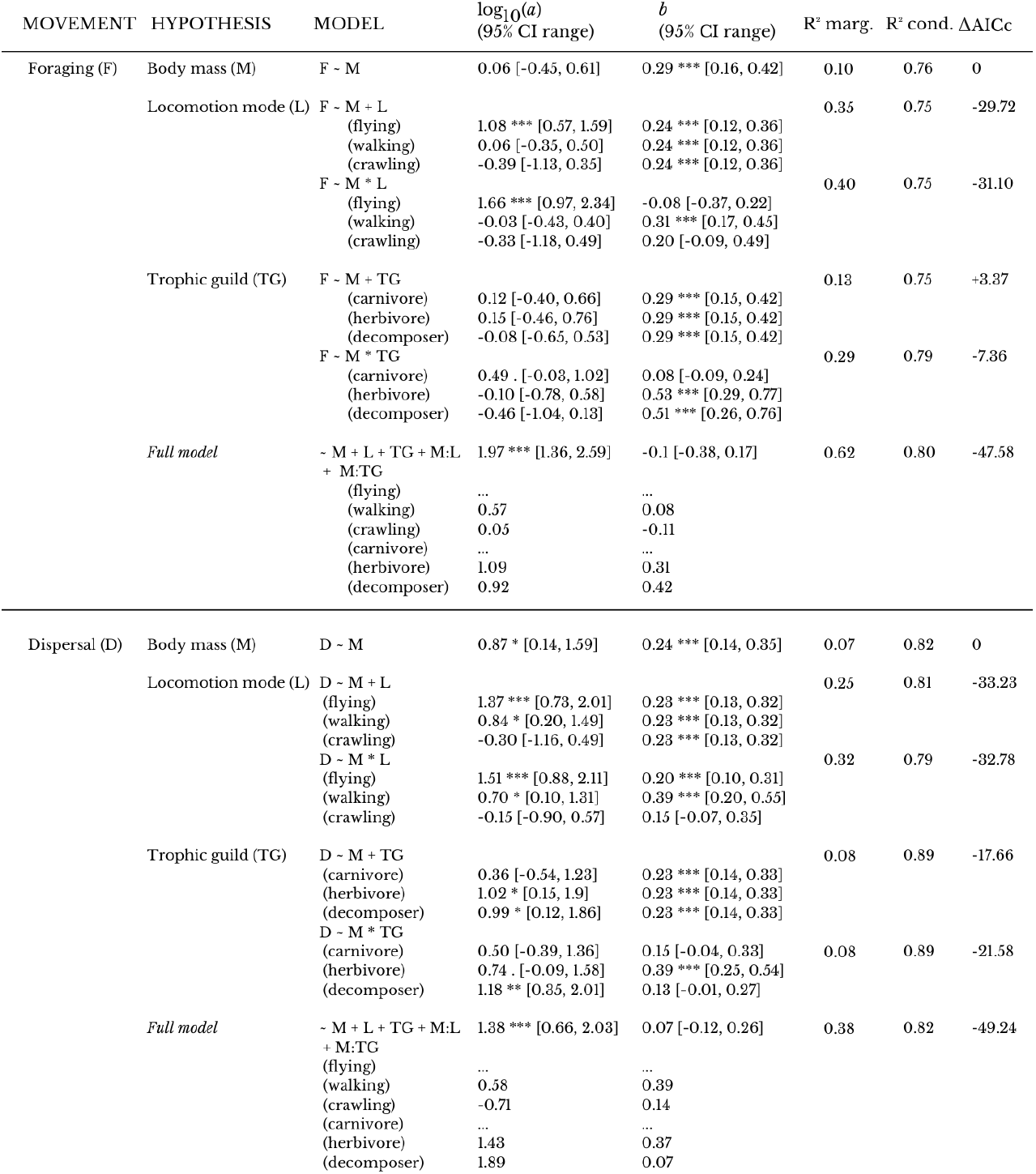
Allometric regression results for linear mixed-effect models. Tracking method and taxonomic order were included in all models as random effects. *log*_*10*_*(a)* : intercept; *b* : slope; R^2^ marg. : marginal R^2^; R^2^ cond. : conditional R^2^; AICc : Akaike information criterion corrected for small sample sizes. Slope and intercept estimates are reported for each factor as absolute values. Significance (p-values) codes : 0 ‘***’ 0.001 ‘**’ 0.01 ‘*’ 0.05 ‘.’ 0.1 ‘ ‘.

**Figure 6.**
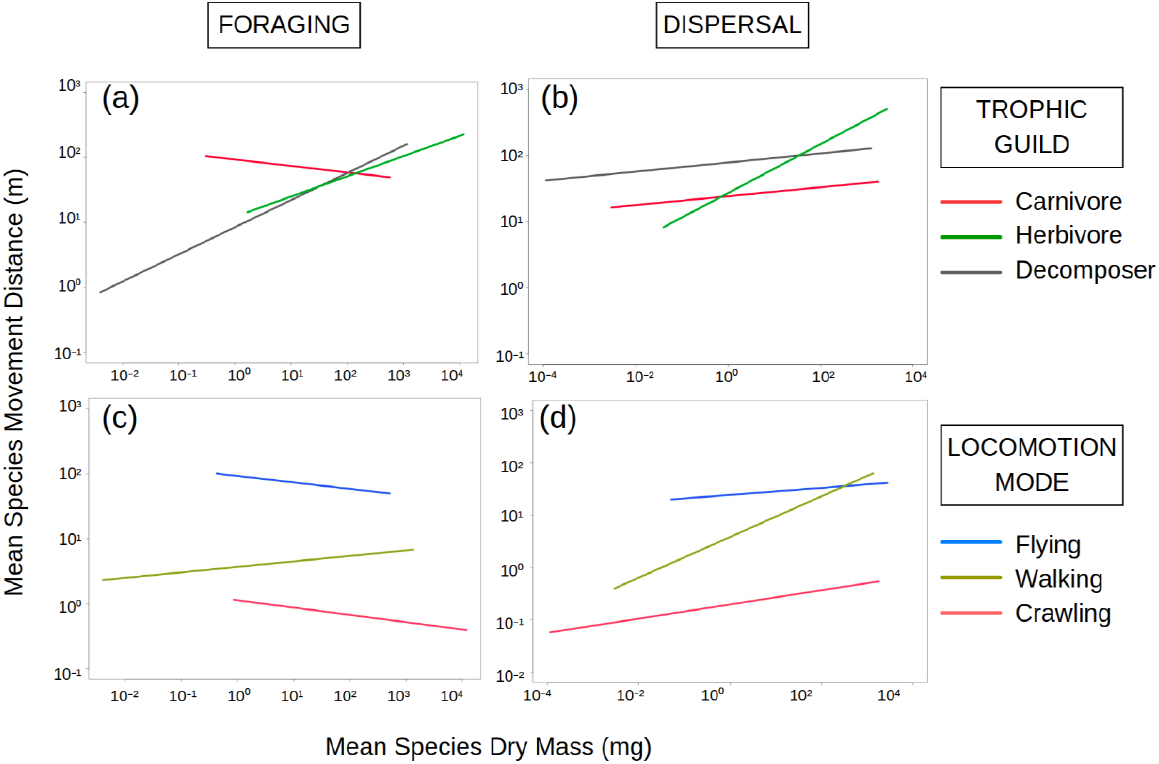
Allometry of space use across trophic guilds and species with similar locomotion strategies, taking taxonomic order and tracking method random effects into account. Regression line coefficients are reported in Table 2 (*Full models* parameters).

We also found a significant interaction effect between locomotion mode and body mass for both dispersal and foraging movements (*ANOVA* : F = 4.538 and 5.202, p = 0.012 and 0.006, respectively) although models with and without this interaction have close AICc values **(Table 2)**. Consistently with our a priori predictions, we found that flying individuals in our database forage further than walkers, themselves moving further than crawlers when controlling for body mass variations (see the blue line that is above the green line that is above the red line in **Figure 6c**). We find the same pattern for the dispersal movements, although we observe a small inversion between flyers and walkers movement distances for the largest organisms **(Figure 6d)**. Flying and crawling individuals show comparable allometric regression slopes, which are negative for foraging and positive for dispersal movements **(Figures 6c and 6d)** when considering full models coefficients **(Table 2)**.

The two environmental factors that were tested, temperature and GPP, only have minor contributions in driving animal movement range for both movement types (partial-*η*^2^ <0.02, **Figure 5**).

## DISCUSSION

We assembled the first global database of movement patterns of terrestrial invertebrates, focusing on active dispersal and foraging movements. Using this database, we documented allometric relationships between body mass and foraging and dispersal movement distances across major terrestrial invertebrate taxonomic orders. We then assessed whether invertebrate movements were driven by the same functional traits as those evidenced for vertebrate taxa. We found that, as for vertebrate species, phylogeny, locomotion strategy and diet are the main drivers of variation in those relationships, since they explain together more than 80% of the total variability in dispersal and foraging movements. This echoes the finding that 80% of the variability in vertebrate home range is explained by locomotion mode, trophic guild, foraging dimension and taxonomy (Tamburello et al. 2015). The relatively low contribution of body mass to the variability of movement distance strongly contrasts with previous results on vertebrates. While 24% and 20% of variability is explained by invertebrates’ body mass for dispersal and foraging movements respectively in simple linear models, these values fall down to 0.07% and 0.10% (marginal R-squared values) when considering taxonomy and tracking method as random effects. In a similar analysis conducted on vertebrate species, Tamburello et al. (2015) evidenced that body mass alone explained up to 44% of home range variability. We identify two main lines of explanation for this discrepancy. First, other morphological traits that do not necessarily correlate with body mass may be better predictors of movement capacities in many invertebrate taxa. In flying invertebrates for example, wing morphology (length, area, elongation) or wing loading (*i*.*e* ratio body mass:wing area) may be more determinant for movement distance than body mass alone (Flockhart et al. 2017). Second, while terrestrial vertebrate movements are measured *in situ* with a small number of available methods, many invertebrate movements can only be studied in experimental systems due to the individuals’ smaller size or underground habitat. The methodological heterogeneity among *in situ* and *ex situ* studies of invertebrates are likely to reinforce the dispersion of data points and to reduce the relative weight of the tested predictors.

Consistently with our predictions, longer dispersal and foraging distances are observed in flying organisms when controlling for body mass, while walking and crawling organisms travel distances that are about one to three orders of magnitude shorter, respectively. We conclude that, as in vertebrate species, invertebrate movement distances correlate with the cost of transport associated with the movement media (Shepard et al. 2013). Our prediction that carnivores would move further than herbivores and decomposers to compensate for lower resource densities is partially supported by our foraging data, but not for dispersal. In the present dataset however, we lack information on potential interaction effects between locomotion mode and trophic guild, due to a strong lack of balance between these two variables in our databases. We dealt with an over-representation of flying individuals in the dispersal database, and of walking individuals in the foraging one, with trophic guilds not being equally represented in both databases (see Table 1 in Supporting information). Still, results of several complementary analyses are globally consistent, reinforcing the robustness of our results. Neither mean temperature of the warmest quarter nor gross primary productivity at the landscape scale appeared to significantly explain the observed interspecific variability in movement distance, although environmental conditions are known to modify the movement behavior of animals in several ways (Johnson et al. 1992). An explanation for this discrepancy is that we tested the influence of global climatic and environmental factors, while individual movement behavior is also driven by weather conditions at finer temporal and spatial scales (*e*.*g*. daily temperature, wind velocity, Knight et al. 2019). Since most studies did not report weather conditions, we were not able to incorporate these environmental variables in our model selection framework.

Our synthesis reveals an over-representation of arthropods in invertebrate movement studies (themselves being dominated by Coleopterans), while extensive data on the space use of annelids, molluscs, nematodes and more generally of the tiniest species are still needed. Tracking methods initially designed for vertebrates, like telemetry or harmonic-radar, have only recently become suitable for studying the movement of the largest invertebrate species (Kissling et al. 2014). The development of innovative tracking methods should improve the spectrum of animals whose movements might be studied in future years (see for example Cointe et al. 2023). For the tiniest invertebrates however, the contribution of active movements to overall displacements is likely to strongly decrease compared to the contribution of passive phoretic movements, especially for dispersal. The exact mechanisms and the spatial extent of phoresy processes remain unclear (Bartlow and Agosta 2021). It would thus be inaccurate and misleading to extrapolate our dispersal data to phoretic animals following our allometric equations.

Our synthesis offers ready-to-use allometric equations to predict terrestrial invertebrate active movements from the sole knowledge of their body mass and a small set of additional functional traits (locomotion mode, diet and body mass). This new information is pivotal for a number of applications, such as the prediction of future species ranges under climate change (Mammola et al. 2021), the design of agroecological landscapes favoring biological control (Haan et al. 2020) or the analysis of connectivity issues for conservation planning (Keeley et al. 2021). More fundamentally, our study also highlights the similarities and differences between vertebrate and invertebrate movements. While we recovered that similar functional traits were driving both vertebrate and invertebrate movements, such as body mass, locomotion mode and diet, the relative influences of these different drivers strongly differ between vertebrate and invertebrate taxa. Although body mass significantly positively correlates with dispersal and foraging distances among the majority of invertebrate orders, its predictive power is clearly lower for invertebrate taxa compared to vertebrate ones. More subtle and taxon-specific approaches might therefore be needed to refine movement inferences from invertebrate traits.

## Acknowledgements

The authors thank the National Research Agency (ANR), the French government IDEX-ISITE initiative 16-IDEX-0001 (CAP 20-25) - CIR 1 - Axis “Fluxes” and Clermont Auvergne Metropole for the funding of this research.

## DATA ACCESSIBILITY STATEMENT

Data will be published on Zenodo after acceptance of this manuscript in a peer-reviewed journal.

## SUPPORTING INFORMATION

**Appendix 1 –** Compiled database

**Appendix 2 –** R script

**Appendix 3 -** Calculation details for the database compilation.

**Appendix 4 -** Full research strategy and additional figures.

## Author contributions

G.A, F.J and J.P conceived the ideas; G.A collected the data, performed the analysis and wrote the successive drafts. F.J, J.M and J.P reviewed the manuscript and proposed improvements.

