## Supplementary information for "Space use of invertebrates in terrestrial habitats: phylogenetic, functional and environmental drivers of interspecific variations"

**SUPPLEMENTARY INFORMATION – Full research strategy and additional figures**

**Research strategy**

We conducted a literature search on the Web of Science and Google Scholar with the following title request :

TI=((invertebrate OR insect OR nematode OR mollusca OR collembola OR diplopoda OR chilopoda OR araneae OR acari OR opiliones OR odonata OR orthoptera OR mantodea OR hemiptera OR diptera OR coleoptera OR lepidoptera OR hymenoptera OR neuroptera OR ant OR spider OR termite OR millipede OR centipede OR earthworm OR slug OR woodlice OR isopoda OR springtail OR blattodea) AND ("space use" OR movement OR "home range" OR territory OR forage) NOT (freshwater OR marine OR aquatic OR benthic OR plankton OR stream)) NOT (WC=(Neurosciences OR "Biochemistry Molecular Biology" OR "Biotechnology Applied Microbiology" OR "Engineering Electrical Electronic" OR "Plant Sciences" OR "Limnology" OR "Pharmacology Pharmacy" OR Telecommunications OR "Computer Science Interdisciplinary Applications" OR "Engineering Biomedical" OR "Fisheries" OR "Horticulture" OR "Toxicology" OR "Food Science Technology" OR "Public Environmental Occupational Health" OR "Meteorology Atmospheric Sciences"))

**Database**

Species in our database were classified either as carnivores, herbivores or decomposers to avoid a multiplication of specific trophic habits that would have small sample sizes and for which we would not have clear *a priori* predictions. A species with a peculiar trophic guild, *Linepithema humile* (May 1868), mainly feeding on honeydew, was considered as a carnivore due to its occasional feeding on small arthropods (Nygard et al. 2008).

**Additional figures**

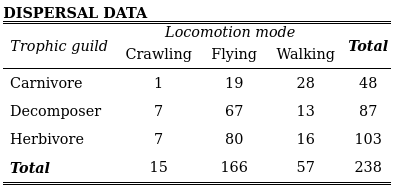

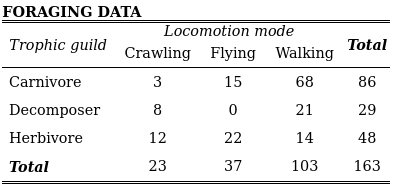

*Table 1. Descriptive statistics of the two datasets. The numbers represent the number of observations, or lines in the databases, corresponding to a given trophic guild and locomotion mode.*

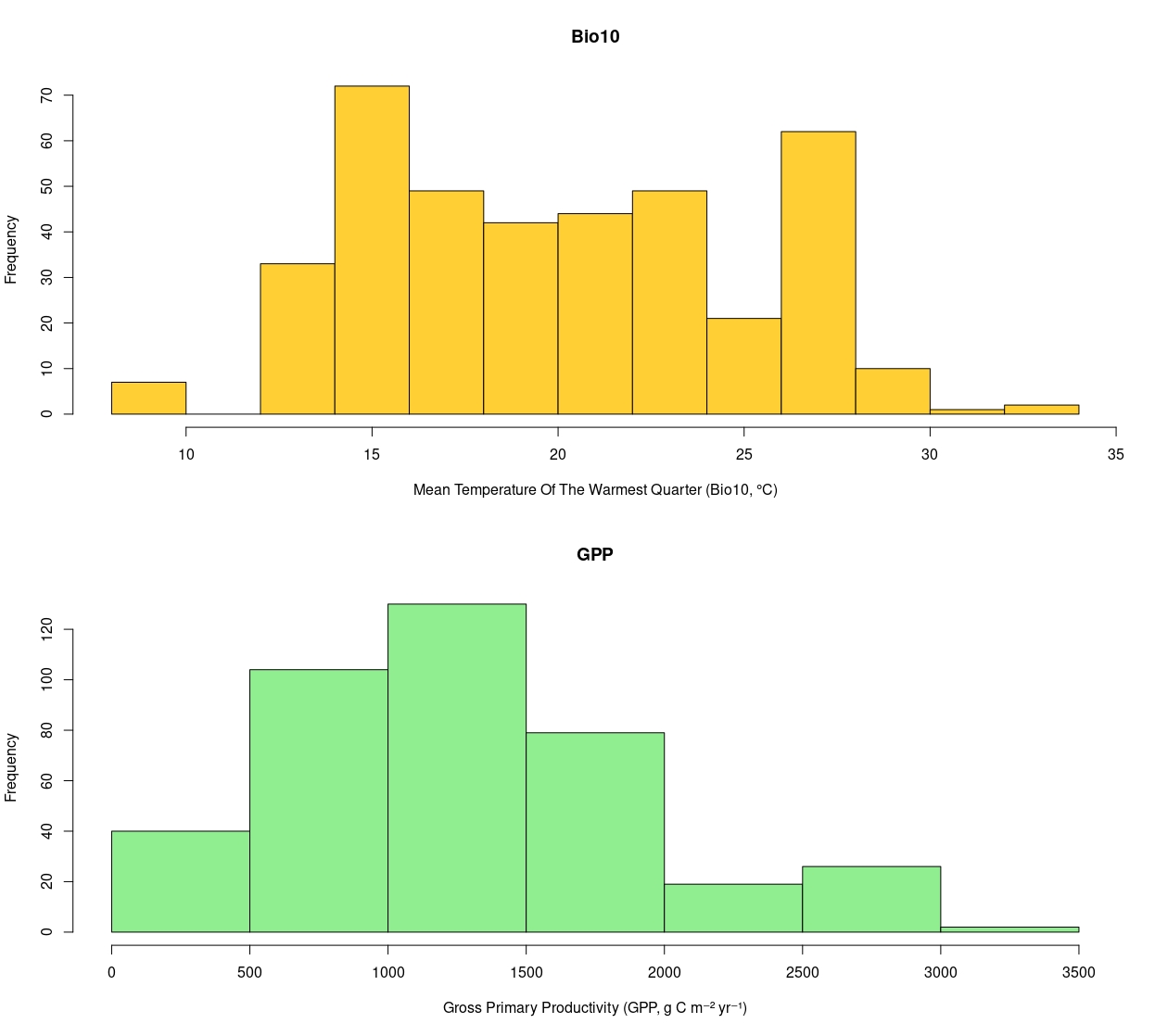

*Figure 1. Range of values for the two environmental variables included in our analysis.*

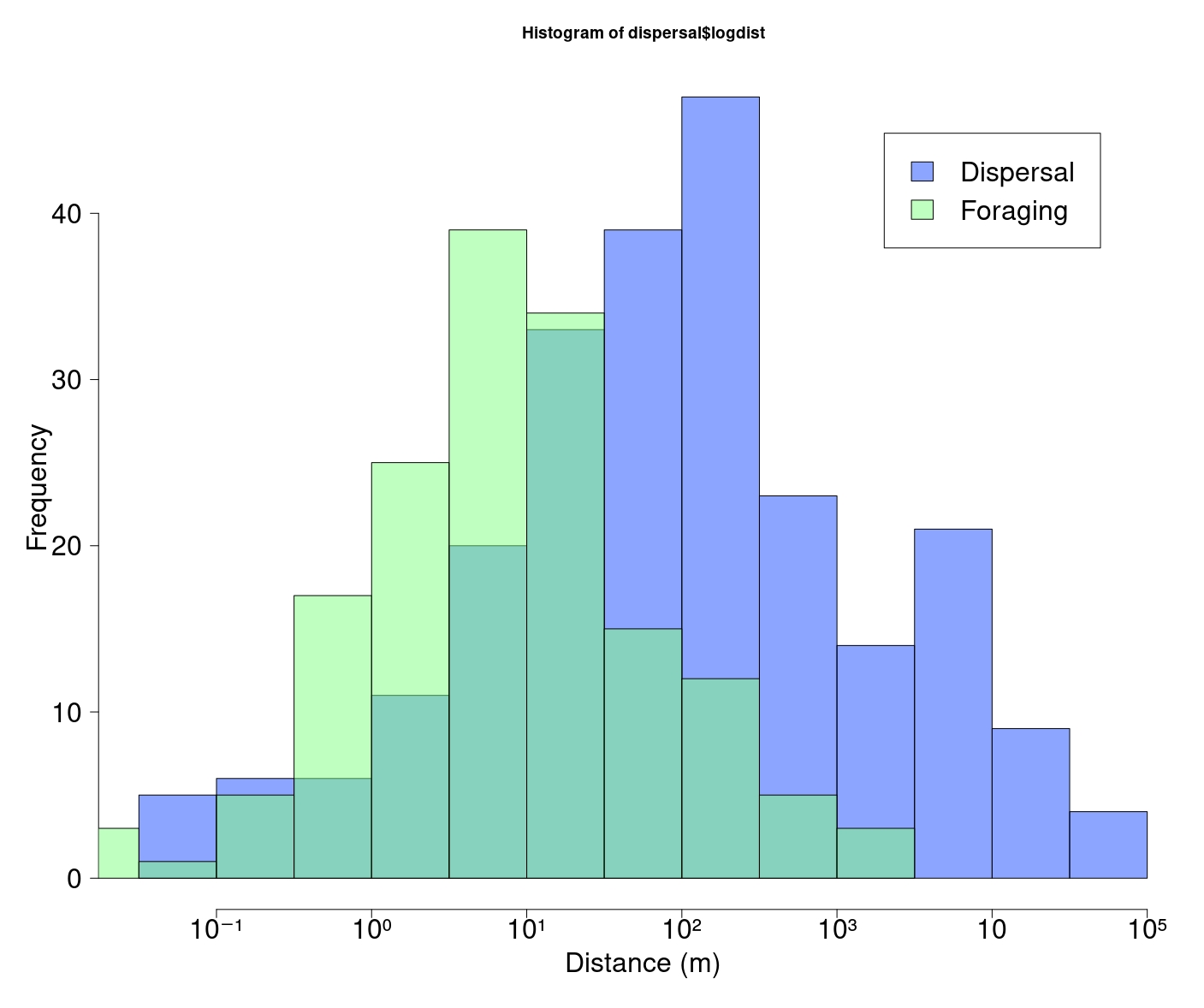

*Figure 2. Distribution of dispersal and foraging distances compiled in the database.*

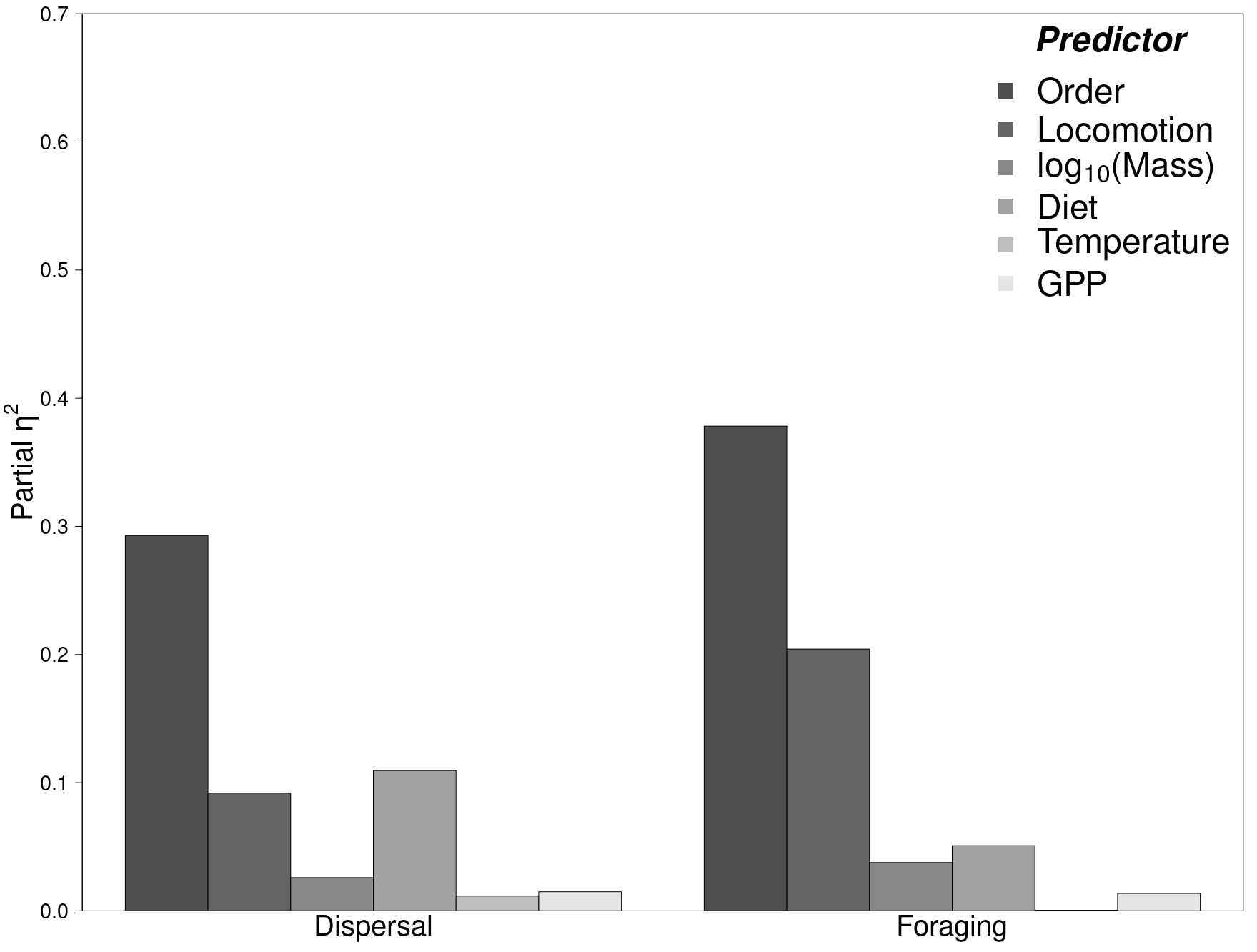

*Figure 2. Effect sizes of taxonomic order, locomotion mode, body dry mass, diet, temperature and GPP as drivers of invertebrate movements. Tracking method is included in the models as random effects. Note that in situ studies only have been considered for this figure to allow the inclusion of the two environmental variables.*
